# NEAT-DNA: A Chemically Accurate, Sequence-Dependent Coarse-Grained Model for Large-Scale DNA Simulations

**DOI:** 10.1101/2025.11.07.687145

**Authors:** Ivan Riveros, Bin Zhang

## Abstract

DNA’s physical properties play a central role in genome organization and regulation, but simulating its behavior across biologically relevant scales remains a major computational challenge. Coarse grained DNA models have enabled faster simulations, yet they often sacrifice chemical accuracy or produce unphysical conformations, limiting their utility for studying genome structure. A key difficulty has been constructing a model that is both efficient enough for large-scale simulations and faithful to the molecular mechanics of DNA. Here we introduce NEAT-DNA, a new coarse-grained DNA model that resolves longstanding limitations in physical realism and parameter optimization. By combining a physically principled energy formulation with a unified training framework that integrates data from both atomistic simulations and experiments, NEAT-DNA accurately reproduces sequence-dependent structure and flexibility while remaining computationally efficient. This approach marks a significant advance over previous models, which either lacked sequence specificity or introduced distortions inconsistent with experimental observations. NEAT-DNA bridges this gap, offering a high-fidelity yet tractable representation of DNA suitable for exploring chromatin folding. More broadly, it provides a foundation for large-scale simulations that couple molecular detail with gene-level chromatin organization, opening new avenues for predictive modeling in structural genomics.

## Introduction

Double-stranded DNA plays a central role in numerous scientific disciplines, including molecular biology, biophysics, and materials science. A key aspect of its functional versatility lies in its ability to adopt a diverse range of conformations, which can be modulated by the underlying nucleotide sequence. This conformational plasticity is fundamental to many biological processes, including gene regulation,^1–3^ chromatin organization,^4–6^ and biomolecular assembly,^7–9^ and is increasingly leveraged in DNA-based nanotechnology.^10^

Computer simulations have emerged as a powerful tool for probing the physical principles underlying DNA conformations and for guiding the rational design of DNA-containing systems. However, the size and complexity of these systems often place them beyond the practical reach of all-atom simulations with explicit solvent. To address this limitation, numerous coarse-grained (CG) models have been developed that eliminate explicit solvent and ions, and represent DNA using a reduced number of interaction sites per nucleotide.^11–27^

Among these, the 3SPN framework and its variants have gained widespread adoption due to their demonstrated accuracy in capturing sequence-dependent mechanical properties of DNA.^14,15,28^ The model has been implemented in molecular dynamics packages such as LAMMPS^29^ and OpenMM,^30,31^ and has enabled the simulation of large-scale assemblies, including oligonucleosomes^32–37^ and protein–DNA complexes,^7,38^ particularly when combined with compatible CG protein force fields.^39,40^

Despite these advances, the computational cost of 3SPN remains nontrivial, limiting its utility in simulations of very large systems such as chromatin segments spanning hundreds of kilobases. To overcome these limitations, several groups have proposed single-bead-per-nucleotide models that offer significantly improved computational efficiency. For instance, the Molecular Renormalization Group (MRG) model developed by the Papoian group^11^ introduced the concept of fan bonds to preserve the local geometry and helical structure of DNA. Initially formulated with explicit ions, MRG was later extended to an implicit electrostatics formulation.^7^ More recently, the Orozco group introduced CGeNArate,^22^ a sequence-dependent single-bead model derived via a bottom-up optimization protocol that integrates atomistic simulations and accounts for sequence-specific mechanical properties.

While these single-bead models are markedly more efficient, their mechanical fidelity becomes compromised under conditions of extreme bending. As we demonstrate below, existing models produce unphysical configurations when DNA is subject to high curvature, such as in nucleosome-bound states. This limits their applicability in chromatin simulations, where such configurations are biologically essential.

In this work, we introduce NEAT-DNA (Nucleosome-ready, Experimentally and Atomically Trained DNA model), a next-generation single-bead CG DNA model specifically designed to maintain physical realism across a broad range of mechanical environments, including those involving tight DNA wrapping around histone proteins. We identify the root cause of the unphysical behavior observed in prior models and present a modified potential energy formulation that resolves this issue. The model is systematically parameterized using a potential contrasting approach based on configurations drawn from explicit-solvent atomistic simulations,^41,42^ and further refined through a reweighting scheme to match experimental measurements of sequence-dependent DNA persistence length.

We show that NEAT-DNA achieves excellent computational efficiency while producing physically realistic conformations across diverse simulation scenarios. The model accurately captures sequence-dependent flexibility and preserves DNA structural integrity under bending stress. We anticipate that NEAT-DNA will enable large-scale, mechanistically faithful simulations of DNA-containing biomolecular assemblies, including chromatin domains^43^ and nuclear condensates.

## Results

### Limitations of MRG-like DNA Models in Capturing Nucleosomal Geometry

DNA undergoes substantial bending stress when wrapped around histone proteins to form nucleosomes, the fundamental repeating units of chromatin. In this highly curved configuration, 150 base pairs of DNA make two superhelical turns around the histone core, deviating markedly from the relaxed geometry of free DNA. To evaluate the performance of existing CG models under such conditions, we simulated nucleosomal DNA using two widely used onebead-per-nucleotide models: the implicit-ion version of MRG^11^ and the CGeNArate model.^22^ Both models offer computational efficiency and share similar energy functions, though CGeNArate additionally incorporates sequence-dependent parameters. Prior work has shown that these models accurately reproduce global properties of long linear and circular DNA molecules.^11,22^ We refer to this family of models collectively as *MRG-like*.

Simulations revealed that MRG-like models produce highly distorted and non-physical geometries when applied to histone-bound DNA. A recurring feature was the tendency of the DNA to adopt square or triangular conformations around the histone core, with long, nearly linear segments interspersed by sharply kinked regions. As shown in Figure 1A, both MRG and CGeNArate produce 3–4 distinct kinks (highlighted in green), separated by extended straight segments. Although CGeNArate exhibited somewhat less severe distortions, it too failed to reproduce the smooth, continuous curvature characteristic of nucleosomal DNA observed in experimental structures.

To quantify this behavior, we introduced a metric termed the *local chain angle* (LCA),^44^ illustrated in Figure 1B. The LCA measures the local bending at each base-paired site along the DNA backbone. As shown in Figure 1C, MRG-like models exhibit an LCA distribution with a pronounced tail toward small angles, consistent with the presence of persistent, localized kinks. These conformations deviate markedly from those observed in crystal structures and atomistic simulations of the nucleosome.^45^

**Figure 1:**
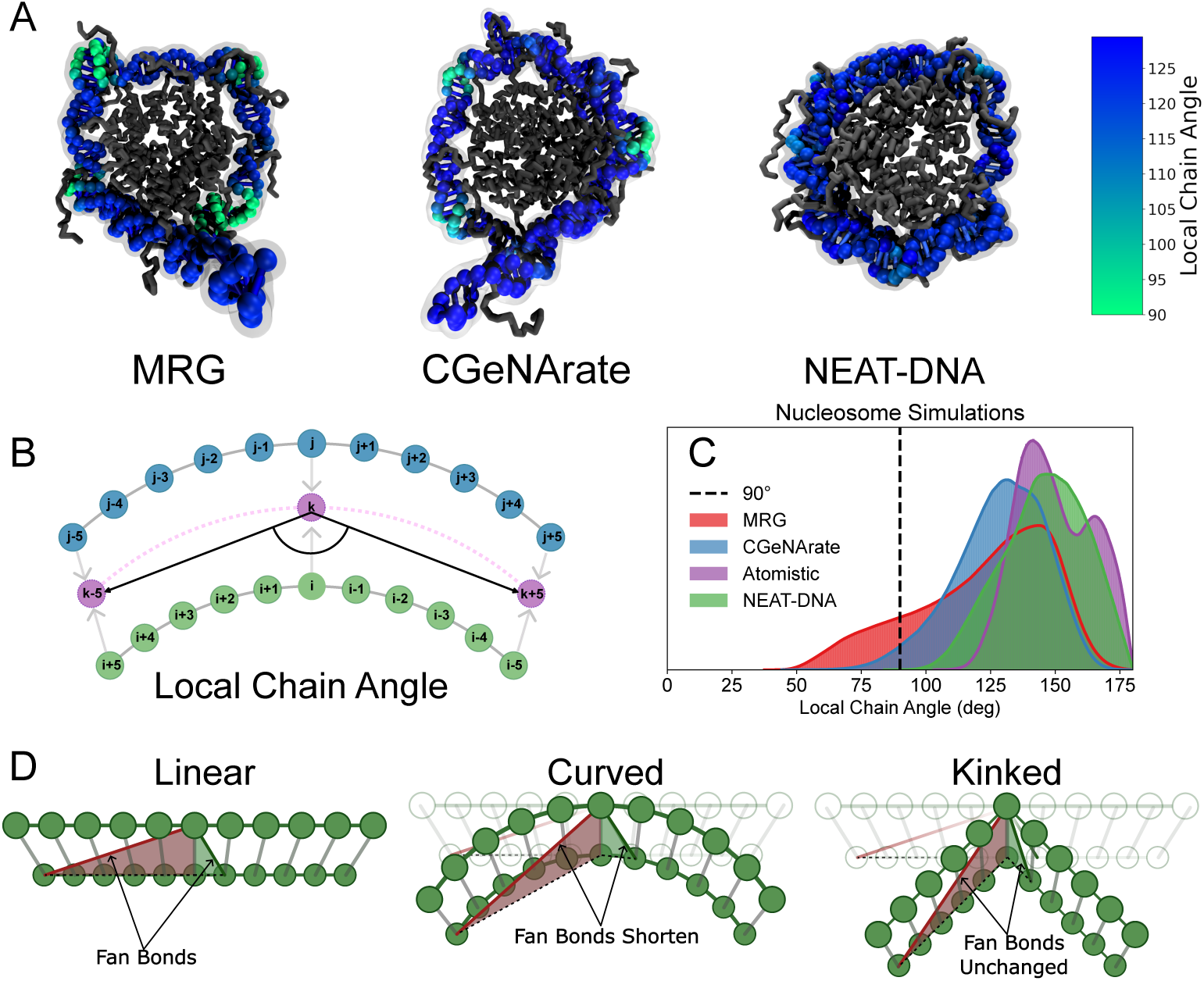
Compared with MRG-like models, NEAT-DNA more accurately reproduces the smooth curvature of nucleosomal DNA by correcting artificial kinks. (A) Representative nucleosome conformations from 10 *µ*s simulations using the MRG, CGeNArate, and NEAT-DNA models, with histones shown in grey and DNA in blue. Regions with distorted geometries, identified by the local chain angle, are highlighted in green. (B) Schematic of the local chain angle (LCA) metric used to quantify local bending or kinks along the DNA backbone. Green and blue beads represent the standard CG sites in both MRG-like and NEAT-DNA models, while violet beads indicate positions used for LCA calculation, obtained by averaging the coordinates of base-paired sites (e.g., defining *k* as the midpoint between *i* and *j*). The LCA at a given base-paired site (*i, j*) is defined by the angle formed by beads *k* − 5, *k*, and *k* + 5. (C) Distributions of local chain angles for each force field compared with atomistic simulations (data from ref. 45). (D) Illustration of how histone-induced bending perturbs the fan bonds in MRG-like models. Long- and short-range fan bonds are shown in red and green, respectively, for linear, uniformly curved, and kinked conformations, demonstrating how kinking preserves equilibrium lengths for most fan bonds.

We hypothesize that the origin of these distortions lies in the design of the *fan bond* potential, which captures long-range cross-strand interactions to preserve DNA helicity (Figure 7C). In both MRG and CGeNArate, fan bonds are implemented using a Class 2 potential (Eq. 6), which is commonly used for covalent bonds due to its steep energetic penalty for deviations from equilibrium. While appropriate for local interactions, this functional form is overly restrictive for non-covalent constraints such as fan bonds.

**Figure 2:**
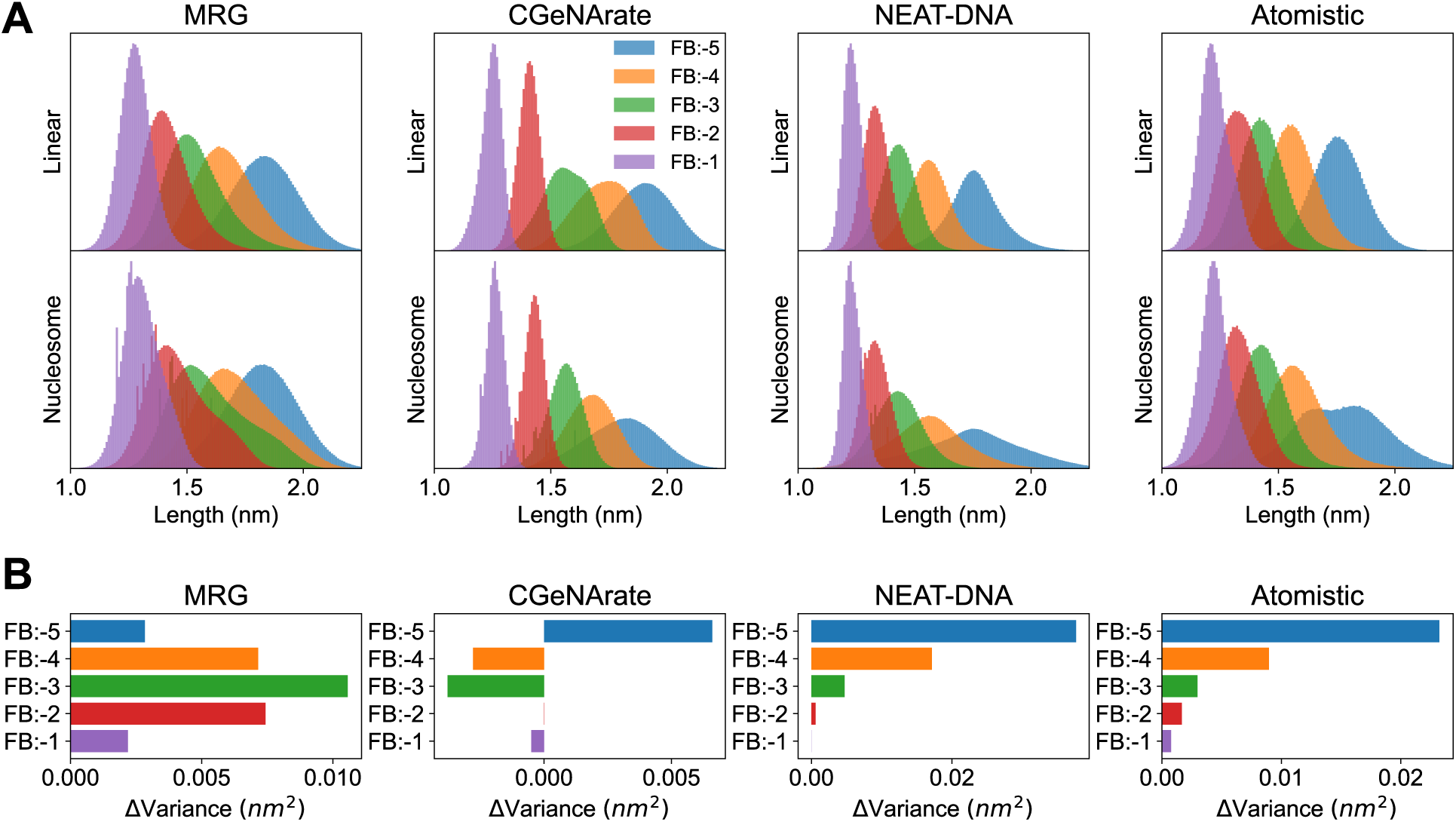
Comparison of fan-bond flexibility between linear and nucleosomal DNA conformations. (A) Distributions of fan-bond lengths in linear and nucleosomal DNA for each coarse-grained model, compared against atomistic references. (B) Quantification of broadening, expressed as the change in variance between the linear and nucleosomal regimes. Atomistic distributions for linear DNA are taken from the miniABC dataset;^41^ nucleosomal data are from ref. 45.

In the nucleosomal context, wrapping DNA around the histone octamer imposes substantial geometric strain, particularly on longer-range fan bonds. This strain leads to significant stretching or compression of these bonds, often driving them far from their equilibrium lengths, an effect visualized in Figure 1D. The steep penalty associated with the Class 2 potential prevents such deviations, causing the system to resolve the resulting frustration by sharply kinking short DNA segments. These short regions absorb the excess strain, while the remainder of the chain remains in a relatively unstressed, linear-like conformation, resulting in the square-shaped geometries shown in Figure 1A.

To verify this hypothesis, we examined the distribution of fan bond distances in simulations of both linear and nucleosomal DNA, and compared these with distributions from atomistic simulations. As shown in Figure 2A, atomistic simulations reveal significant broadening of the longest-range fan bond (FB:-5) in the nucleosomal context relative to linear DNA, consistent with the expected stretching and compression during histone wrapping. Moreover, Figure 2B shows that this broadening trend extends across all fan bonds, with the magnitude of deviation decreasing for shorter-range terms.

While the CGeNArate model does capture some broadening of FB:-5, it underestimates the extent of this effect, and fails to reproduce the expected trend for other fan bonds. This discrepancy reinforces the conclusion that the use of overly rigid fan bond potentials in MRG-like models leads to non-physical geometries in nucleosomal simulations, and highlights the need for more flexible formulations to accurately model stressed DNA conformations.

### Systematic Optimization of NEAT-DNA Using Atomistic and Experimental Data

We set out to develop a CG DNA model capable of accurately capturing nucleosome-bound configurations, motivated by the limitations of existing MRG-like models. Our goal was to address these shortcomings while retaining the strengths of previous models. We term the resulting model *NEAT-DNA*, which stands for **N**ucleosome-ready, **E**xperimentally and **A**tomically **T**rained DNA model.

As described in the *Methods* section, a central design improvement of NEAT-DNA is the reformulation of the energy function governing interactions between nucleotides on opposite strands. To resolve the issues associated with fan bonds in MRG-like models, we replace the Class 2 potential with a Gaussian potential (Eq. 12). This formulation is inspired by the Gō-type potentials commonly used in CG protein folding models^39,46^ and was recently adopted in RNA modeling.^47^ Unlike the Class 2 potential, which imposes an increasingly large penalty as the bond deviates from its equilibrium length, the Gaussian potential has a finite energy cost for bond rupture. This modification enables the model to accommodate the geometric frustration experienced by DNA in highly curved configurations such as those in nucleosomes.

**Figure 3:**
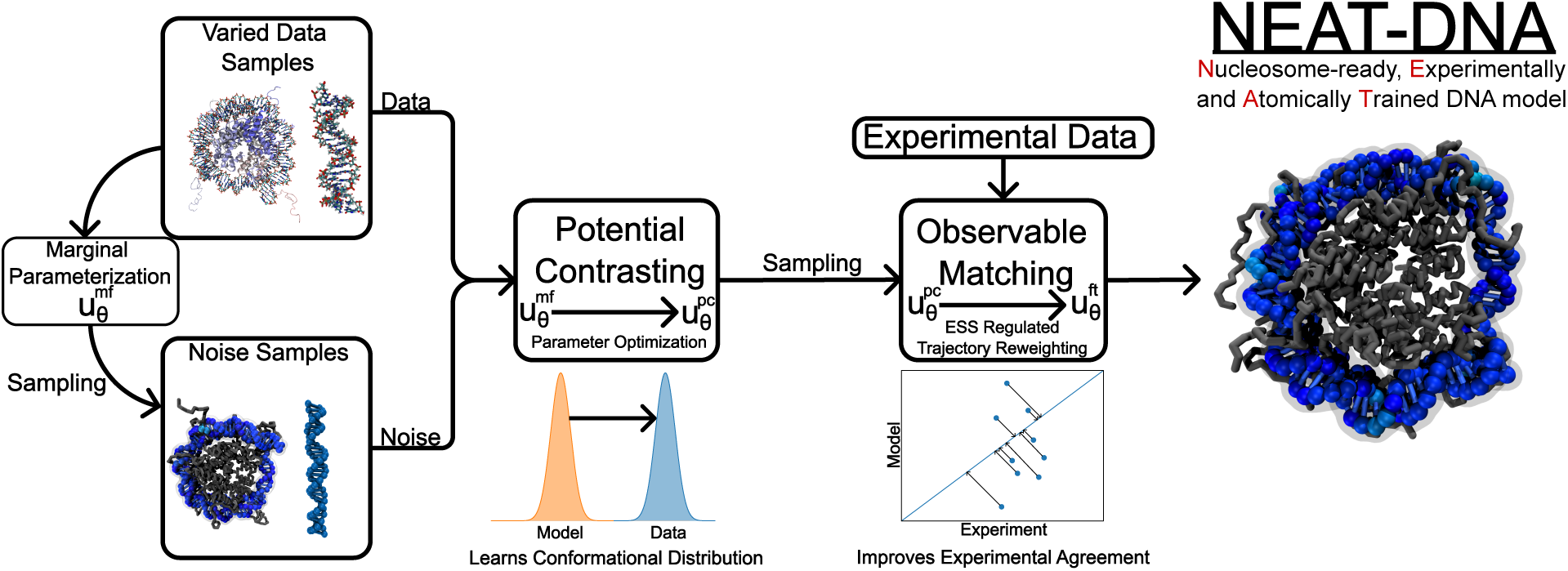
Schematic of the parameterization workflow for NEAT-DNA. Model parameters are first optimized using potential contrasting, which aligns the energy function with the high-dimensional configurational distributions present in the training datasets. Following this, the parameters are fine-tuned to reproduce experimental observables, such as sequence-dependent DNA persistence lengths.

In addition to distance-dependent interactions, the fan bond potential also incorporates angular and dihedral terms (Eq. 12) to improve the representation of B-form DNA geometry.^14,15,28,48–51^ These terms modulate the strength of the distance-based Gaussian potential, effectively softening its contribution when angular or torsional strain is present. This design reduces the energetic penalty for distance deviations caused by DNA bending, and allows the model to maintain physically realistic structures under stress. Similar angular dependencies defined at the trimer level were introduced to encode sequence specificity in the model (Eq. 3).

Model optimization followed a two-stage scheme analogous to the pretraining–fine-tuning paradigm used in machine-learning force fields^52–60^ (see *Methods* and Figure 3). In the pretraining stage, parameters were adjusted using atomistic ensembles from the miniABC dataset^41^ and nucleosome simulations.^45^ Potential contrasting^61^ was employed to ensure that the model reproduced the configurational distributions present in the atomistic reference data. Subsequently, parameters were fine-tuned using an ensemble reweighting procedure to reproduce experimentally determined persistence lengths across diverse DNA sequences.

By combining atomistic and experimental information, NEAT-DNA bridges the gap between bottom-up^61–64^ and top-down modeling paradigms.^38,65–68^ The bottom-up stage captures detailed molecular interactions and sequence-dependent flexibility from atomistic data, while the incorporation of experimental observables compensates for limitations of the underlying atomistic force fields. This integrative approach allows NEAT-DNA to retain near-atomistic accuracy while achieving experimentally consistent mechanical properties.

### Validation of NEAT-DNA Across Structural, Sequence, and Experimental Benchmarks

Before assessing sequence-dependent and global structural accuracy, we examined whether the new energy formulation in NEAT-DNA resolves the geometric artifacts observed in MRG-like models. As shown in Figures 1A and 1C, nucleosome simulations with NEAT-DNA no longer exhibit the distorted “square” or “kinked” geometries characteristic of MRG and CGeNArate. The probability distribution of the local chain angle closely matches that obtained from atomistic nucleosome simulations, indicating a more physically realistic backbone conformation. Furthermore, NEAT-DNA preserves short-range bonded interactions while allowing longer-range fan bonds to break under bending stress, consistent with atomistic behavior (Figure 2).

We next assessed NEAT-DNA’s ability to reproduce atomistic configurations across both local and global structural scales. At the local level, we compared bond length and angle distributions for various trimers against atomistic references from the miniABC dataset. As shown in Figure 4A and Figure S1, NEAT-DNA faithfully reproduces sequence-dependent distributions with accuracy comparable to, or in many cases exceeding, that of CGeNArate, as quantified by the Kullback-Leibler (KL) divergence. At the global level, we evaluated configurational fidelity by comparing root-mean-square deviation (RMSD) distributions between atomistic and CG ensembles. As shown in Figure 4B, NEAT-DNA accurately reproduces the atomistic RMSD profiles across multiple sequences, demonstrating that the model captures the overall configurational ensemble rather than merely matching local statistics.

**Figure 4:**
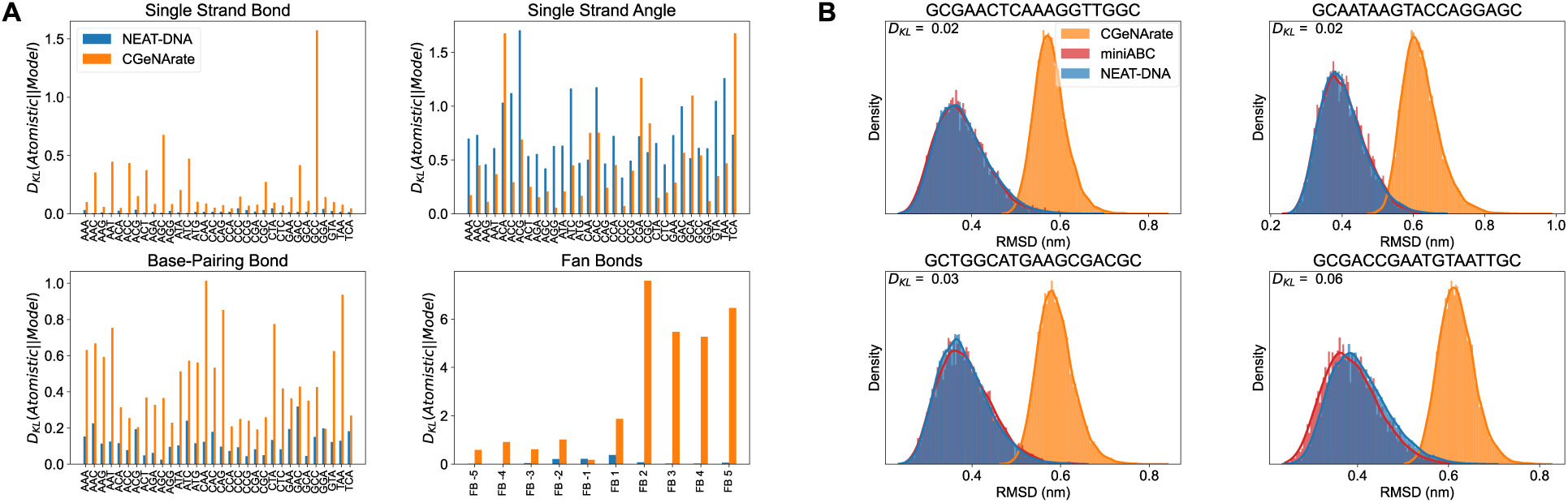
NEAT-DNA accurately reproduces sequence-dependent configurational distributions from atomistic simulations. (A) Kullback–Leibler (KL) divergence comparing sequence-specific distributions obtained from coarse-grained models against atomistic references in the miniABC dataset. Shown are single-strand bond (top left), single-strand angle (top right), and base-pairing bond (bottom left) distributions across 32 representative trimers (reduced from 64 by symmetry). The bottom-right panel reports KL divergence for fan-bond distributions relative to the miniABC dataset. (B) Examples of the two best (top row) and two worst (bottom row) performing trimers based on KL divergence, showing root-mean-square deviation (RMSD) distributions relative to the mean atomistic structure. Distributions from miniABC trajectories and from coarse-grained models are overlaid for comparison.

The agreement is especially noteworthy given that, after pre-training on atomistic reference data, NEAT-DNA was further fine-tuned against experimental observables. This outcome underscores the effectiveness of potential contrasting in capturing higher-order structural correlations and demonstrates that fine-tuning, aided by the effective sample-size regularizer,^52^ preserves consistency with atomistic statistics by constraining parameter updates within statistically supported regions of the atomistic ensemble.

Consistent with the fine-tuning objective, NEAT-DNA also accurately reproduces known sequence-dependent mechanical properties. As shown in Figure 5A and B, the model predicts persistence lengths with a Spearman’s rank correlation of *ρ* = 0.7 and an RMSE of 3.7nm relative to experimental measurements,^69,70^ demonstrating the effectiveness of the top-down parameterization. In comparison, CGeNArate, which lacks top-down calibration, achieves a moderate correlation of *ρ* = 0.6 and relatively high RMSE of 16.7nm.

**Figure 5:**
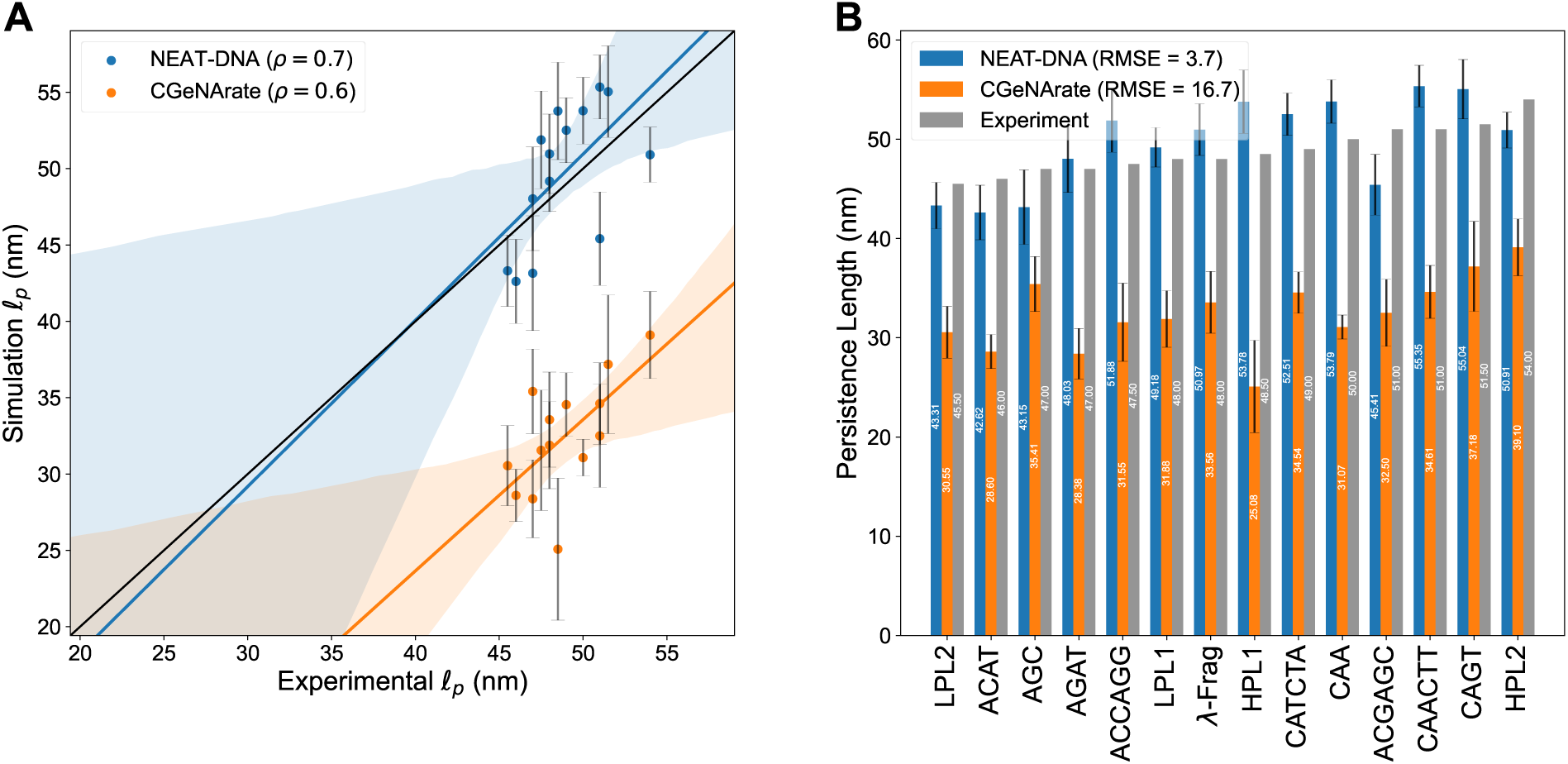
NEAT-DNA accurately reproduces experimental persistence length values across diverse sequences. (A) Correlation between experimentally measured persistence lengths and predictions from NEAT-DNA and CGeNArate. Lines of best fit with shaded confidence intervals are provided as visual guides. Numerical values of all persistence lengths are listed in the Supporting Information. (B) Sequence-resolved comparison of predicted persistence lengths from both models. Error bars denote 95 % confidence intervals estimated by block averaging. Note that the *λ*-Frag sequence^70^ was not utilized in training.

Finally, we benchmarked the computational performance of NEAT-DNA. Because it employs a one-bead-per-nucleotide resolution similar to MRG and CGeNArate, NEAT-DNA achieves comparable simulation speeds. Using our OpenMM implementation, a 200bp duplex (400 particles) runs at approximately 35*µ*s/day on an NVIDIA RTX2080Ti GPU, whereas 3SPN.2C, a higher-resolution but nucleosome, capable model—reaches only 5*µ*s/day under equivalent conditions. The performance gap indeed widens with longer sequences (Figure 6).

**Figure 6:**
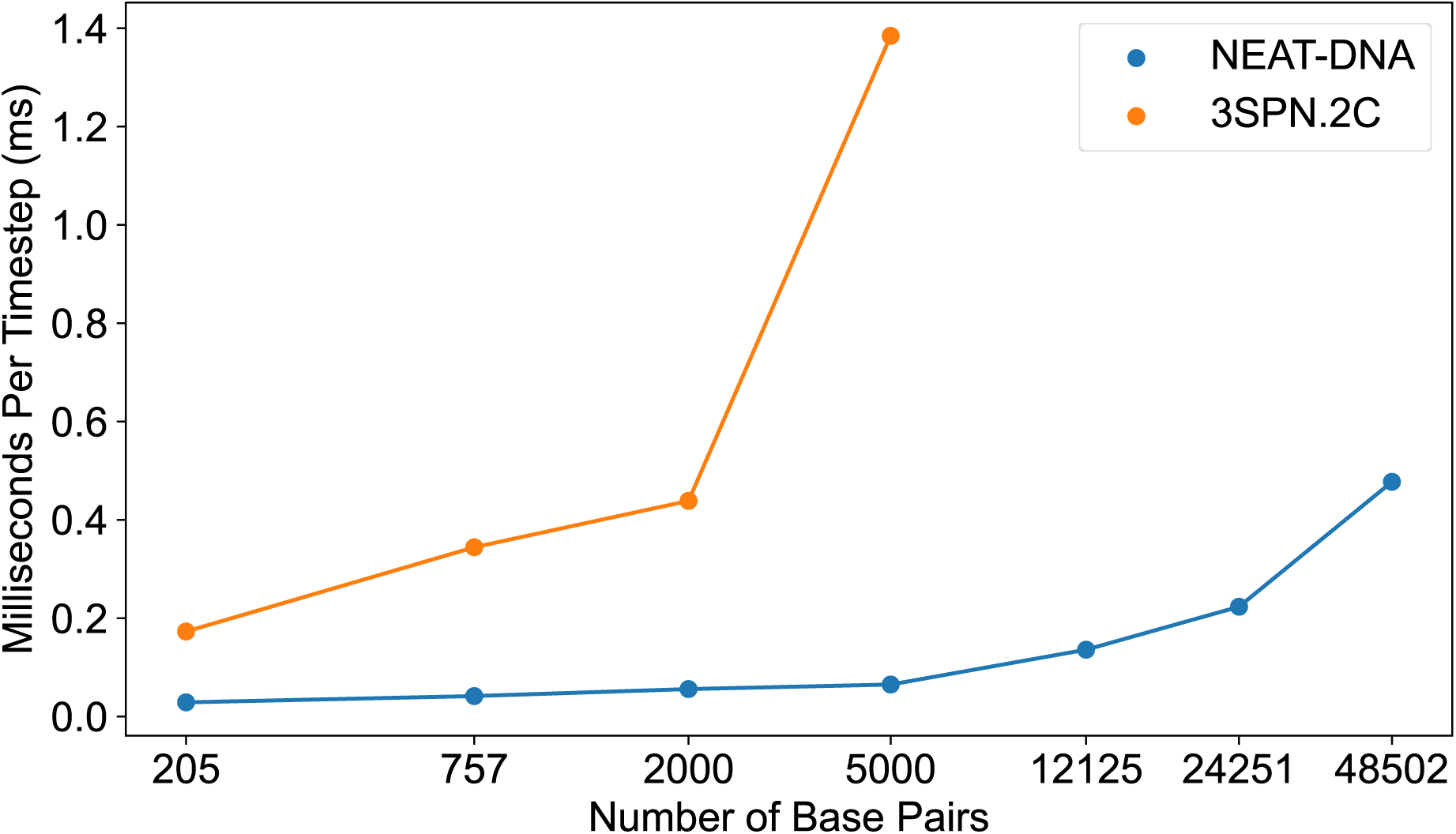
Scaling performance of NEAT-DNA and 3SPN.2C with increasing DNA length. Average simulation time per integration step as a function of DNA length is plotted for NEAT-DNA and 3SPN.2C.^15^ All benchmarks were performed on NVIDIA RTX 2080 Ti GPUs using the OpenABC^71^ implementation of each force field within OpenMM.^30^ Data for 3SPN.2C are unavailable beyond 12,125 bp due to excessive memory requirements.

## Conclusions and Discussion

We have presented NEAT-DNA, a next-generation CG DNA model designed to capture sequence-dependent mechanics and nucleosome-scale configurations with high physical fidelity and computational efficiency. A central innovation is the reformulation of the fan bond potential, which replaces the overly rigid Class 2 potential used in prior models with a flexible Gaussian-based form that accommodates DNA bending without inducing unphysical kinks. This modification eliminates the square-like conformations observed in MRG-like models and yields backbone geometries that closely resemble those in atomistic nucleosome simulations.

NEAT-DNA was parameterized through a multi-stage strategy that combines atomistic simulations and experimental data within a unified optimization framework. Potential contrasting was used to match atomistic structural ensembles across a wide range of DNA conformations, while differentiable trajectory reweighting allowed fine-tuning of sequence-specific mechanical properties to reproduce experimental persistence lengths. This hybrid training approach led to a model that not only preserves near-atomistic accuracy but also generalizes well across structural regimes inaccessible to either atomistic simulation or bottom-up models alone.

Compared to CGeNArate, NEAT-DNA adopts a more parsimonious representation of sequence specificity. While CGeNArate uses overlapping tetramers, our trimer-based scheme avoids double-counting and simplifies force decomposition, thereby facilitating model interpretability and downstream integration with multimodal systems.

We further note that the comprehensive optimization protocol employed here, spanning marginal parameter fitting, potential contrasting, and observable-matching fine-tuning, has not been previously applied to CG force field development at this level of integration. The ability to reconcile atomistic and experimental benchmarks across local and global structural scales suggests that this framework may be broadly applicable to the construction of other multiscale models.

Importantly, NEAT-DNA achieves simulation speeds comparable to other one-bead-per-nucleotide models while resolving the geometric pathologies that have hindered their application to chromatin-scale systems. When combined with similarly CG representations of histones and other nuclear proteins, NEAT-DNA opens the door to efficient, mechanistically faithful simulations of large-scale chromatin organization and dynamics.

## Methods

### Energy function of NEAT-DNA

NEAT-DNA employs a coarse-grained model of double-stranded DNA, representing each nucleotide as a single bead. The potential energy associated with a given configuration is defined as:

**Figure 7:**
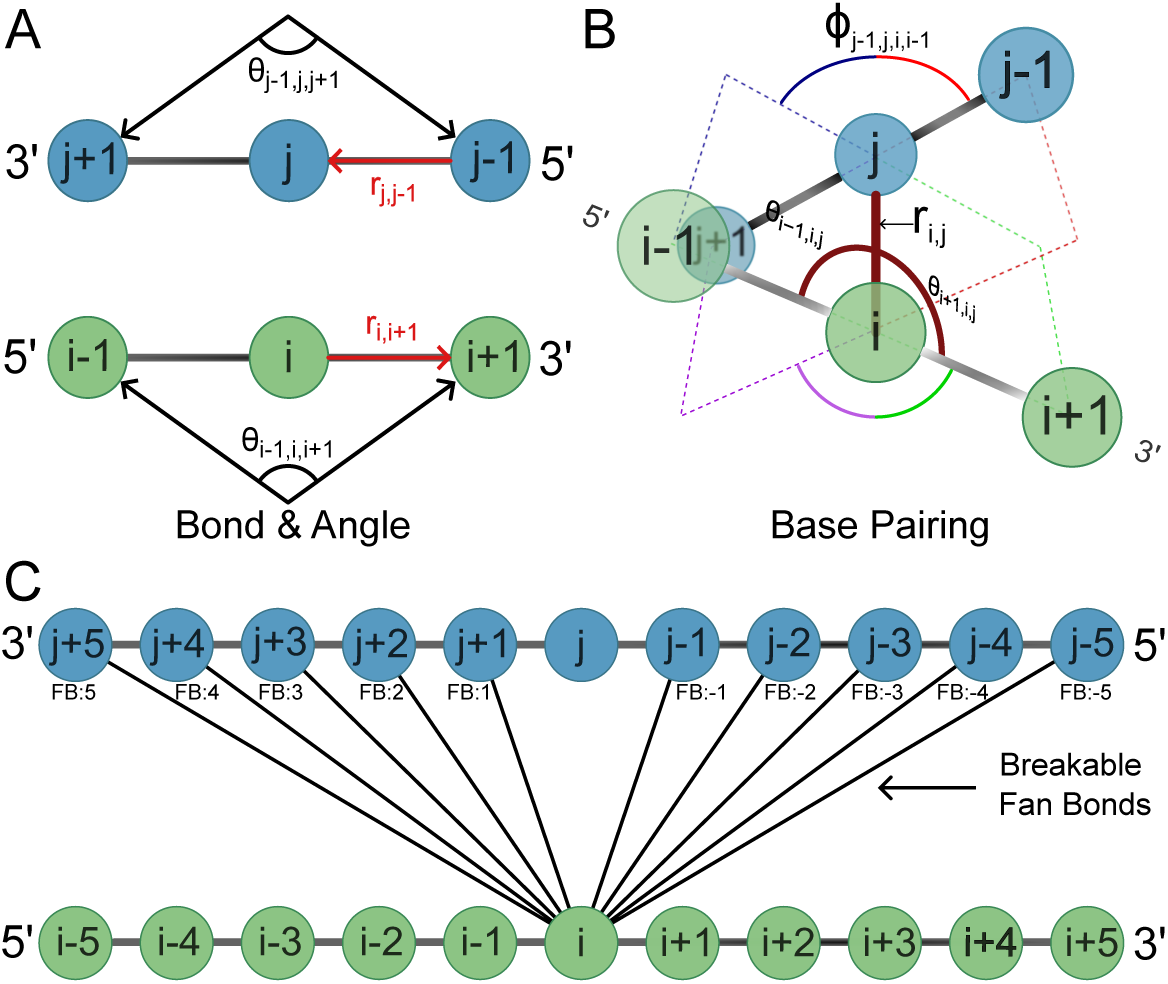
Visualization of force field components. (A) Schematic of the bonded and angular terms comprising 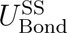 and 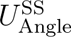. (B) Illustration of the base-pairing and cross-strand interactions included in *U*_Pairing_. The dihedral angles are defined by the blue/red and purple/green planes, respectively. Together with the cross-strand angle terms and the bond distance, these interactions describe the geometry of paired bases. (C) Visualization of cross-strand interactions contributing to *U*_BFB_, corresponding to fan bonds in MRG-like models.

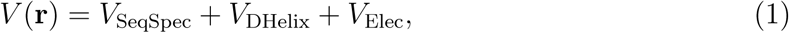

where **r** denotes the coordinates of the nucleotides on both strands. The three terms on the right-hand side correspond, respectively, to: a sequence-specific potential that preserves local base pairing, *V*_SeqSpec_; a term that encodes the helical geometry of DNA, including major and minor grooves, *V*_DHelix_; and a screened electrostatic interaction term, *V*_Elec_. Detailed definitions of each term are provided below.

**Sequence-specific potential.** To encode sequence-dependent behavior, we introduce the following potential for each base-paired nucleotide pair (*i, j*):

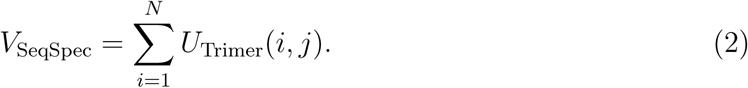

The trimer potential *U*_Trimer_(*i, j*) depends on the positions of *i* and *j*, as well as the adjacent upstream and downstream nucleotides on both strands. For terminal nucleotides that lack the required neighboring bases, the corresponding energy terms are omitted.

Our formulation is similar in spirit to the sequence-dependent force field introduced in CGeNArate,^22^ which also incorporates local sequence context to modulate bonded and angular interactions. However, whereas CGeNArate uses overlapping tetramers to define local sequence environments, our approach assigns each base-paired nucleotide pair to a single trimer context centered on the pair. This avoids the double counting of interactions that can arise in overlapping tetramer schemes and ensures a unique energy contribution for each local sequence motif.

The trimer potential *U*_Trimer_(*i, j*) comprises three components:

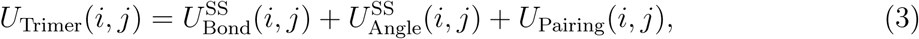

where 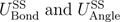 act on nucleotides within the same strand (Figure 7A) and are given by

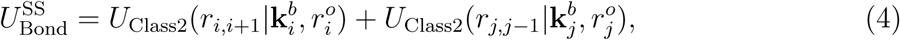

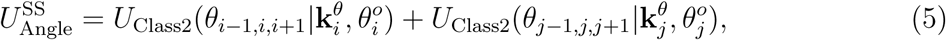

with the Class 2 potential defined as

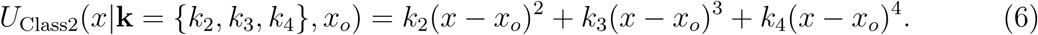

The base-pairing term *U*_Pairing_(*i, j*) includes cross-strand bond, angle, and dihedral (Figure 7B) interactions:

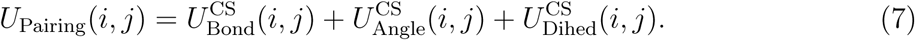

The cross-strand bond term is defined as

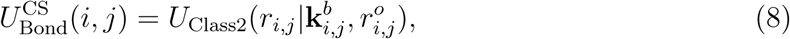

representing the interaction between nucleotides that form a base pair.

The angular component includes four angles defined among the six nucleotides within the trimer:

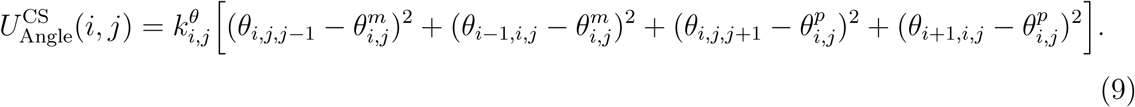

The dihedral term includes two torsional angles involving the six nucleotides in the trimer:

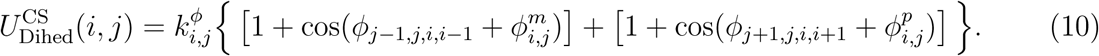

The spring constants 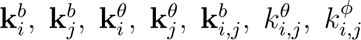 and the equilibrium values 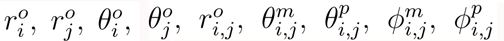 are parameters that depend on the identity of the 64 possible Watson–Crick base-paired DNA trimers.

**Double helix potential.** The helical structure of DNA is stabilized by a series of noncovalent, sequence-independent interactions captured by the double helix potential, defined as

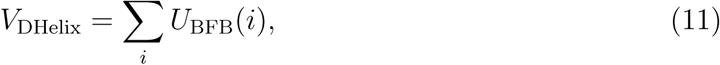

where *U*_BFB_(*i*) represents a breakable fan bond (BFB) interaction centered at nucleotide *i*:

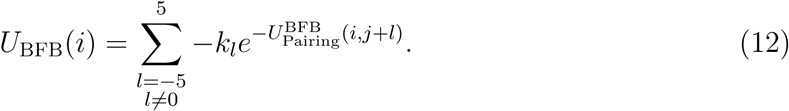

Here, *j* denotes the nucleotide base-paired with *i*, and *k_l_*is a force constant associated with the cross-strand interaction between *i* and its *l*-th neighbor on the complementary strand. The interaction energy 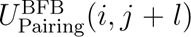 follows a similar functional form as *U*_Pairing_(*i, j*) in Eq. 7, but with a simplified bonded term:

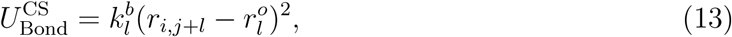

where 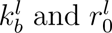 are the spring constant and equilibrium distance, respectively, for the interaction between *i* and *j* + *l*.

**Electrostatic potential.** Electrostatic interactions between nucleotides are modeled using the Debye-Hückel potential:

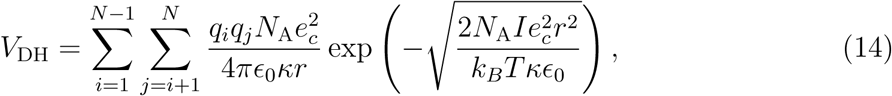

where *q_i_* and *q_j_* are the partial charges assigned to nucleotides *i* and *j*, respectively; *N*_A_ is Avogadro’s number; *e_c_* is the elementary charge; *κ* is the dielectric constant of water; *ε*_0_ is the vacuum permittivity; *I* is the ionic strength of the solution; *T* is the absolute temperature; *k_B_* is Boltzmann’s constant; and *r* is the distance between the two charges. To account for counterion condensation, we set *q_i_*= −0.6, following the theoretical framework of Manning^72^ and the parametrization introduced by Lequieu et al. ^73^ .

### Parameterization of NEAT-DNA

We parameterized the NEAT-DNA model in three stages (Figure 3), marginal fitting (mf), potential contrasting (pc), and fine-tuning (ft). Technical details are provided in the *Supporting Information*.

**Marginal fitting to atomistic structural distributions.** As a first step, we constructed an initial force field by independently fitting each component of the potential energy function to marginal distributions derived from atomistic simulations in the miniABC dataset.^41,74^ These components include the bonded and angular terms for single strands, 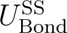 (Eq.4) and 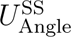 (Eq.5); the cross-strand bonded, angular, and dihedral terms for base pairing, 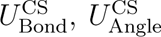 (Eq.9), and 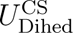 (Eq.10); and the cross-strand interactions in the breakable fan bond potential (Eq. 12).

To construct reference distributions, we extracted trimer segments from the atomistic simulations and grouped together segments of the same type. By assuming symmetry between each trimer and its base-paired complement (e.g., AGC was treated as equivalent to GCT), we reduced the dataset to 32 unique representative trimers. For each trimer type, we computed the distributions of bond lengths, angles, and dihedral angles, which were then used to parameterize the corresponding components of the potential energy function. We refer to the resulting initial model as the *NEAT-mf* force field, to distinguish it from the fully optimized version described in the following section.

**Global optimization with diverse DNA structural ensembles.** To jointly optimize all model parameters and leverage a broader spectrum of DNA conformations, we applied the potential contrasting method,^61^ a scalable framework for high-dimensional parameter spaces and large structural datasets.

The training data comprised approximately 80,000 configurations drawn from three sources:

- MiniABC dataset: 5,000 atomistic configurations from each of the thirteen 18-mer duplexes,^41,74^ providing equilibrium B-form DNA structures under physiological conditions.
- Nucleosome simulations: 10,000 configurations from explicit-solvent atomistic MD of nucleosome-bound DNA,^45^ capturing highly bent conformations stabilized by histone-DNA contacts. These structures are inaccessible to the linear miniABC sequences but reflect biologically relevant curvature.
- 3SPN simulations: 5,000 configurations from a long-timescale CG simulation of a 198-bp linear duplex, using the explicit ion 3SPN model.^28,75^ These data introduce dynamic transitions between straight and moderately bent states, complementing the static curvature in nucleosomes and the near-linear geometries in miniABC.

Together, these datasets span a range of curvatures and contexts, from unconstrained thermal fluctuations to protein-induced bending, providing a comprehensive structural ensemble for global parameter refinement.

Potential contrasting updates the model parameters by optimizing an energy function that discriminates between real (training) and noise configurations. The noise ensemble was constructed from molecular dynamics trajectories generated with the NEAT-mf model:

- MiniABC sequences: 13 independent 10 *µ*s simulations (one per sequence).
- 3SPN sequence: a 50 *µ*s simulation of the 198-bp duplex.
- Nucleosome sequences: two 10 *µ*s simulations corresponding to PDB IDs 1KX5 and 3LZ0.

From each trajectory, 5,000 configurations were uniformly sampled to construct the noise distribution.

All optimization procedures were implemented in PyTorch^76^ using the Adam optimizer,^77^ with NEAT-mf parameters as initialization. The resulting globally optimized model is referred to as NEAT-pc.

**Fine-tuning with Experimental Data.** To improve the model’s agreement with experimentally measured sequence-dependent persistence lengths, we fine-tuned the NEAT-*pc* force field using persistence length data for 13 DNA sequences obtained from DNA cyclization experiments by Geggier and Vologodskii ^69^ .

We employed differentiable trajectory reweighting^78^ to estimate how small perturbations in the energy function parameters affect an observable, without requiring new simulations. Instead, expectations under a modified potential are computed from a single reference trajectory via reweighting. The reweighted estimator for an observable O is given by

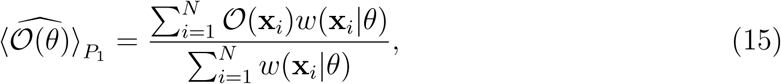

where 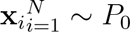 are samples from the reference ensemble, and the reweighting factor is

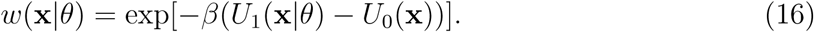

Here, *U*_0_(**x**) denotes the potential energy that generated *P*_0_ (i.e., the original NEAT-*pc* model), *U*_1_(**x**|*θ*) is the perturbed potential under parameters *θ*, and *β* is the inverse thermal energy. Because Eq. 15 is differentiable with respect to *θ*, it can be directly optimized to minimize the discrepancy between predicted and experimental observables, in this case, the persistence length (see the *Supporting Information* for more details).

Following Rufa et al. ^52^, we included an effective sample size (ESS) regularizer to maintain sufficient overlap between the reference (*P*_0_) and reweighted (*P*_1_) ensembles:

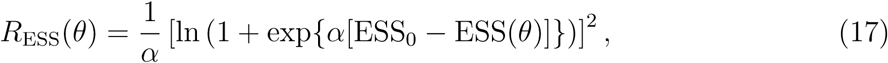

where *α* and ESS_0_ control the strength and offset of the penalty. We set *α* = 100 and ESS_0_ = 750, equal to the values utilized by the Chodera group.^52^ The effective sample size is defined as

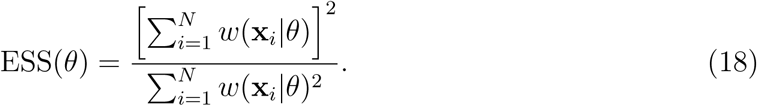

The final loss function for observable matching is then

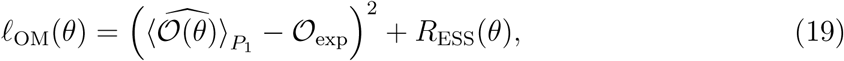

where 𝒪_exp_ is the experimentally determined persistence length for each sequence.

### Molecular dynamics simulation details

NEAT-DNA was implemented within the OpenABC simulation framework,^71^ enabling GPU-accelerated CG molecular dynamics using OpenMM.^30^ All simulations were performed in the canonical (NVT) ensemble at 300K and an ionic strength of 150mM. Dynamics were propagated using the Langevin Middle integrator with a 10 fs timestep, and configurations were saved every 2 ns. For consistent benchmarking, the implicit-ion MRG-DNA^11^ and CGeNArate^22^ models were also implemented in OpenABC.

**Linear DNA simulations.** We performed DNA-only simulations for three sequence sets: (i) the miniABC dataset, (ii) sequences with experimentally measured persistence lengths, and (iii) the nucleosomal DNA sequence (PDB ID: 1KX5). Simulations of the miniABC and nucleosomal sequences were run for 10 *µ*s per sequence, while those in the experimental validation set were extended to 150 *µ*s. Initial configurations were generated by coarse-graining atomistic models constructed with X3DNA,^79^ where each nucleotide was represented by the centroid of its atomic coordinates.

**Nucleosome simulations.** Nucleosome systems containing 147 base pairs of DNA wrapped around a histone octamer were constructed from the crystal structure (PDB ID: 1KX5)^80^ and converted to CG resolution in OpenABC, using one bead per amino acid and per nucleotide. The SMOG force field^39^ was applied to preserve the native conformation of the histone octamer. Protein-DNA interactions were modeled using a Debye-Hückel potential for electrostatics and a Lennard-Jones like potential for excluded-volume effects, following previous studies.^7^ To maintain the wrapped DNA geometry, each nucleotide carried a charge of −1 in the electrostatic calculations. The 5 base pairs centered at the dyad axis, along with the structured regions of the histone octamer, were constrained as a rigid body to prevent nucleosome sliding. Each nucleosome simulation was run for 10 *µ*s.

## Acknowledgement

This work was supported by the National Institutes of Health (Grant R35GM133580) and the National Science Foundation (MCB-2042362). I.R. was the recipient of the National Science Foundation Graduate Research Fellowship. This work used Bridges-2 at PSC through allocation BIO240299 from the Advanced Cyberinfrastructure Coordination Ecosystem: Services & Support (ACCESS) program, which is supported by U.S. National Science Foundation grants #2138259, #2138286, #2138307, #2137603, and #2138296.

## Competing interests

The authors declare that they have no competing interests.

## Data and materials availability

NEAT-DNA is implemented into the open source software package OpenABC. Tutorials for setting up simulations of NEAT-DNA can be found on the GitHub.

## Notes

### Competing Interest Statement

The authors have declared no competing interest.

