## Supporting Information for "NEAT-DNA: A Chemically Accurate, Sequence-Dependent Coarse-Grained Model for Large-Scale DNA Simulations"

### Contents

**Model Parameterization** **S-3**

Marginal . . . . . S-3

Potential contrasting . . . . . S-4

Persistence Length Observable Matching . . . . . S-5

**Supplemental Figures** **S-7**

**Supplemental Tables** **S-8**

**References** **S-9**

### Model Parameterization

#### Marginal

Fitting was performed by converting the miniABC generated distribution associated with the potential term of interest, for example the bond length distribution, to an un-normalized probability distribution via binning. This un-normalized distribution was used to fit the individual potentials by minimizing the following objective:

$$\ell_{\text{marginal}}(\theta) = \left\langle \left( \exp(-\beta U(x|\theta)) - \tilde{P}_R(x) \right)^2 \right\rangle_{P_R}, \tag{S1}$$

where $\tilde{P}_R$ is the un-normalized reference probability distribution, $\theta$ are the parameters of the potential term, and sub-scripted angle brackets, $\langle \cdot \rangle_{P_R}$, denote an expectation over the reference distribution $P_R$. Class 2 potentials additionally had constraints applied to ensure physically-realistic behavior.

Specifically, we ensured the Class 2 potentials could only have a single minima by utilizing the cubic discriminant of the Class 2's derivative. The cubic discriminant for a cubic polynomial $ax^3 + bx^2 + cx + d$ is as follows:

$$\Delta = 18abcd - 4b^3d + b^2c^2 - 4ac^3 - 27a^2d^2 \tag{S2}$$

A useful property of the cubic discriminant is that when $\Delta \leq 0$, it is guaranteed that the cubic polynomial has a single root. Thus, we can use this as a constraint for the Class 2 potential's derivative:

$$\frac{d}{dx}U_{Class2} = 2k_2(x - x_0) + 3k_3(x - x_0)^2 + 4k_4(x - x_0)^3 \tag{S3}$$

$$\Delta_{dU_{Class2}} = 36k_3^2k_2^2 - 128k_4k_2^3 \tag{S4}$$

Solving this equation for $k_3$ when $\Delta_{dU_{Class2}} \leq 0$ gives the following inequality:

$$-\frac{4}{3}\sqrt{2k_4k_3} \leq k_3 \leq \frac{4}{3}\sqrt{2k_4k_3} \tag{S5}$$

By ensuring that $k_2, k_4 > 0$, we can then define the following constraint for $k_3$ to guarantee the potential has a single minima:

$$k_3 = \frac{4}{3}\tanh(k_3')\sqrt{2k_4k_2}, \tag{S6}$$

where $k_3'$ is an internal, hidden value for optimization only. Utilizing this method (as opposed to fixing $k_3$ to 0) allows for a single minima without losing the ability to model skew-symmetric and heavy-tailed potentials.

#### Potential contrasting

The objective function of Potential Contrasting is given by:

$$\begin{aligned}\ell_{PC}(\theta, \Delta F) = &\left\langle \ln\left(\frac{1}{1+\nu\exp\left(-(U_N - U_\theta + \Delta F)\right)}\right)\right\rangle_{\text{Data}} + \\ &\nu\left\langle \ln\left(\frac{1}{1+\nu^{-1}\exp\left(-(U_\theta - U_N - \Delta F)\right)}\right)\right\rangle_{\text{Noise}},\end{aligned} \tag{S7}$$

where $\theta$ are the parameters associated with potential energy function $U_\theta$, $\langle\cdot\rangle_{\text{Noise}}$ is the distribution associated with known potential energy function $U_N$, $\langle\cdot\rangle_{\text{Data}}$ is the target data distribution, in our case the miniABC samples, $\nu$ is the ratio between the number of noise samples and number of data samples, and $\Delta F$ is a learned free energy difference between the noise and learned potentials. Note that at no point during optimization is any sampling of the learned distribution necessary, allowing the optimization process to be efficiently completed in one-shot, differing from other parameterization methods.[S1]

To further regularize parameter changes, we utilized the expected sample size regularizer ($\mathrm{R}_{\mathrm{ESS}}(\theta)$), details in the main article (Eq. 17), with hyper-parameters set to $\alpha = 0.01$ and

$\mathrm{ESS}_0 = 50$ and NEAT-mf as $U_0(\mathbf{x})$ in Eq. 16.

#### Persistence Length Observable Matching

As discussed in the main text, our objective was to use an observable-matching fine-tuning scheme to optimize NEAT-pc against experimentally measured quantities. In this case, the observable of interest is the DNA persistence length ($\ell_p$).

Traditionally, $\ell_p$ is estimated by fitting the correlation function of the DNA chain to an exponential decay,

$$\left\langle \hat{t}(0) \cdot \hat{t}(s) \right\rangle = e^{-s/\ell_p}, \tag{S8}$$

where $\hat{t}(s)$ is the unit tangent vector at contour position $s$. However, because this approach requires arbitrarily choosing the range of $s$ used for fitting, the resulting estimates can be unreliable, especially at small $s$, where the exponential dependence amplifies statistical noise.

To obtain a more robust measure, we instead determined $\ell_p$ using the mean-squared end-to-end distance ($R^2$) of the chain, which can be related to $\ell_p$ through the analytical expression derived from the worm-like chain model:[S2]

$$\left\langle R^2 \right\rangle = 2\ell_p L_c \left[ 1 - \frac{\ell_p}{L_c} \left( 1 - e^{-L_c/\ell_p} \right) \right], \tag{S9}$$

where $L_c$ is the contour length of the DNA. This formulation directly links the experimentally measured persistence length to the simulated distribution of $R^2$, avoiding the need for arbitrary fitting choices and providing a more stable numerical observable for reweighting.

Accordingly, given experimental $\ell_p$ values, we reweighted each ensemble based on its squared end-to-end distance, which serves as a simple yet physically meaningful constraint. For this procedure, we used persistence length measurements from Geggier and Vologodskii[S3] for 13 DNA sequences. Simulations with the optimized NEAT-pc model were performed for these sequences to generate ensembles for reweighting. When comparing with experimental data, Eq. S9 was solved numerically to compute $\ell_p$ from the simulated end-to-end distance distributions.

### Supplemental Figures

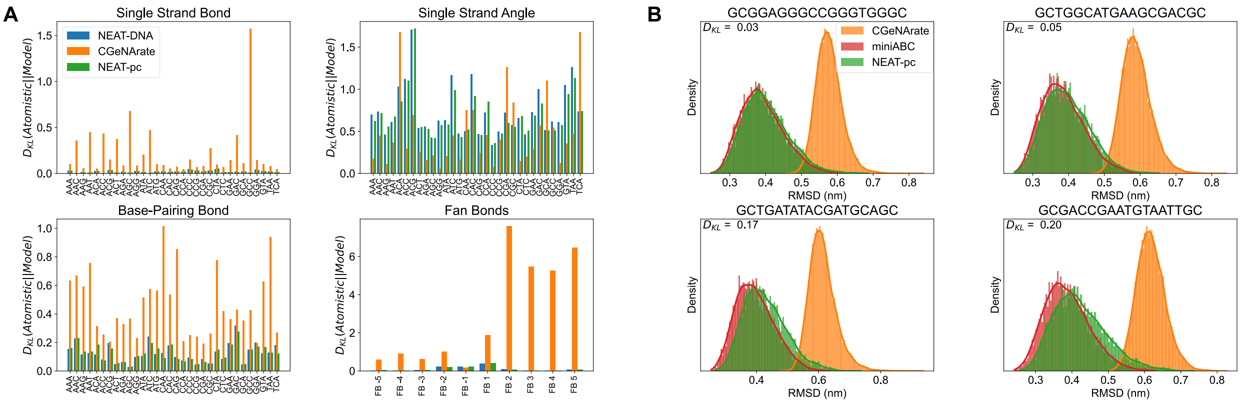


Figure S1: **NEAT-DNA accurately reproduces sequence-dependent configurational distributions from atomistic simulations.** The plots are similar to those presented in Figure 4 of the main text, with the addition of results from the NEAT-pc model.

### Supplemental Tables

Table S1: Comparison of persistence length predictions from NEAT-DNA and CGeNArate with experimental measurements for all 14 tested sequences. Simulation results are reported using both the exponential fitting method (Eq. S8) and the end-to-end distance formulation (Eq. S9). All values are in nanometers (nm). (*) Note that the $\lambda$-Frag sequence was not included in the training set.

| | | $\langle R^2 \rangle$ Based $\ell_p$ | | Exponential Fit Based $\ell_p$ | |
|---|---|---|---|---|---|
| Sequence | Experiment | NEAT-DNA | CGeNArate | NEAT-DNA | CGeNArate |
| LPL2 | 45.5 | $43.3 \pm 2.3$ | $30.6 \pm 2.6$ | $50.6 \pm 3.8$ | $39.9 \pm 5.3$ |
| ACAT | 46.0 | $42.6 \pm 2.8$ | $28.6 \pm 1.7$ | $51.0 \pm 5.3$ | $37.0 \pm 3.6$ |
| AGC | 47.0 | $43.2 \pm 3.8$ | $35.4 \pm 2.8$ | $52.3 \pm 6.7$ | $46.2 \pm 5.7$ |
| AGAT | 47.0 | $48.0 \pm 3.4$ | $28.4 \pm 2.6$ | $57.1 \pm 5.9$ | $34.9 \pm 4.8$ |
| ACCAGG | 47.5 | $51.9 \pm 3.2$ | $31.6 \pm 3.9$ | $63.2 \pm 6.0$ | $37.8 \pm 6.7$ |
| LPL1 | 48.0 | $49.2 \pm 2.0$ | $31.9 \pm 2.9$ | $59.8 \pm 3.5$ | $39.9 \pm 5.4$ |
| HPL1 | 48.5 | $53.8 \pm 3.2$ | $25.1 \pm 4.7$ | $65.4 \pm 5.4$ | $27.5 \pm 6.6$ |
| CATCTA | 49.0 | $52.5 \pm 2.1$ | $34.5 \pm 2.1$ | $65.1 \pm 4.1$ | $47.0 \pm 4.7$ |
| CAA | 50.0 | $53.8 \pm 2.2$ | $31.1 \pm 1.2$ | $66.3 \pm 3.7$ | $39.5 \pm 2.4$ |
| ACGAGC | 51.0 | $45.4 \pm 3.1$ | $32.5 \pm 3.4$ | $55.4 \pm 5.5$ | $41.2 \pm 6.3$ |
| CAACTT | 51.0 | $55.4 \pm 2.1$ | $34.6 \pm 2.7$ | $69.1 \pm 3.9$ | $45.3 \pm 5.7$ |
| CAGT | 51.5 | $55.0 \pm 3.0$ | $37.2 \pm 4.5$ | $67.6 \pm 5.5$ | $46.0 \pm 8.0$ |
| HPL2 | 54.0 | $50.9 \pm 1.8$ | $39.1 \pm 2.9$ | $60.9 \pm 3.0$ | $51.2 \pm 6.0$ |
| $\lambda$-Frag* | 48.0 | $51.0 \pm 2.6$ | $33.6 \pm 3.1$ | $61.4 \pm 4.6$ | $43.5 \pm 6.5$ |
| Spearman's $\rho$ | - | 0.7 | 0.6 | 0.7 | 0.5 |
| RMSE | - | 3.7 | 16.7 | 12.6 | 9.1 |
